# Conformational and Thermodynamic Differences Underlying Wild-type and Mutant ENL YEATS Domain Specificity for Epigenetic Marks

**DOI:** 10.1101/2022.07.14.500085

**Authors:** Lokesh Baweja, Jeff Wereszczynski

## Abstract

Histone post-translational modifications (PTMs) are interpreted by multiple reader domains and proteins to regulate gene expression. The ENL YEATS domain is a prototypical PTM reader that recognizes multiple lysine acetylation marks on the histone H3 tails as a way of recruiting chromatin remodellers. Two ENL YEATS mutations have been identified which have been linked with leukemia, Wilms tumor, and other forms of cancer and result in either an insertion or deletion of residues in the loop connecting beta sheets distant from the protein active site. *In vitro* experiments have shown that these mutations modulate the selectivity of YEATS domains for various lysine acetylation marks, although different experiments have provided contrasting views on the ability of the insertion and deletion mutants to discern specific PTMs. Here, we have performed multiple molecular dynamics simulations of wild-type, insertion, and deletion mutant YEATS domains free from and in complex with two PTM peptides: one that is acetylated at K9 of H3 and the other that is acetylated at residue K27 of H3. Results show that these two peptides have distinct flexibilities and binding energetics when bound to YEATS domains, and that these properties are affected by interactions with residues within and outside of the peptide consensus motif. Furthermore, these properties are modulated by the YEATS insertion and deletion mutants, which results in disparate binding effects in these systems. Together, these results suggest that only the partial exposure of histone tails is sufficient in the context of nucleosomes for YEATS-mediated recognition of acetylation marks on histone tails. They also caution against the over-interpretation of results obtained from experiments on reader domain-histone peptide binding in isolation and not in the full-length nucleosome context.

## Introduction

Chromatin reader proteins process epigenetic information by recognizing post-translational modifications (PTMs) on the histone tails.^1^ Eleven nineteen leukemia (ENL) binds with acetylated chromatin regions through its YEATS domain (named for the first discovered members of the family: Yaf9, ENL, AF9, Taf14, Sas5) ^2,3^ and recruits super elongation complex that regulates transcription of ENL target genes through RNA polymerase II.^2,4^ Oncogenic transformations in acute myeloid leukemia are linked with acylation reading of ENL, and inhibition of the YEATS domain via small molecules is suggested as a promising therapy to treat leukaemia.^5–8^

YEATS adopts an immunoglobulin-like topology with an elongated β-sandwich core with four loops (L1, L4, L6, and L8) connecting the β-strands (Figure 1). The narrow hydrophobic binding site is formed by the three loops (L1, L4, and L6) and lined by histidine, serine, and tyrosine.^9^ YEATS binds to Nε-acetylated lysine (Kac) in histone tails at positions K9 and K27, and in turn, stabilizes the transcription machinery for the expression of target genes.^3,10,11^ To date, eight mutations have been discovered in ENL protein causing insertions or deletions in the L8 region of the YEATS domain.^12,13^ Among the known ENL mutants, insertion (p.117_118insNHL) and deletion (p.111_113NPP*>*K) mutations have been found at a high frequency within diseased patients and nearly 5 percent of people having Wilms tumor.^12,14^

**Figure 1.**
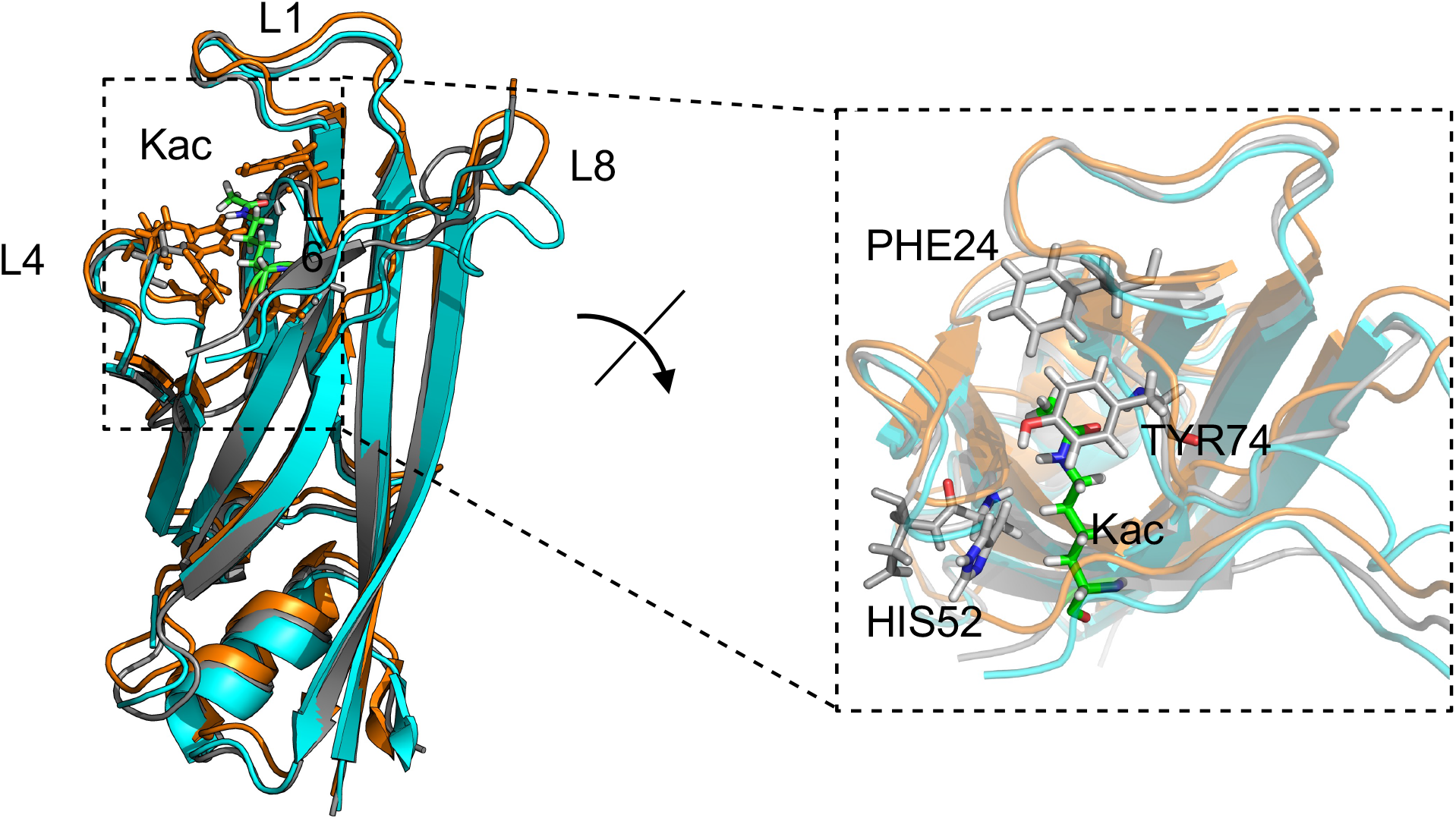
Cartoon representation of wild-type, deletion and insertion mutants of the ENL YEATS domain in complex with full-length H3K9ac peptide. The wild-type YEATS domain is shown in orange, the deletion mutant in gray, and the insertion mutant in cyan. A zoomed-in view of the Kac pocket is provided with the active residues displayed in stick representation.

*In vitro* experiments performed using isolated histone peptide mimics (H3K9ac and H3K27ac) have shown that these insertion and deletions mutants bind H3K9ac and H3K27ac peptides with varied thermodynamics, ^12,15^ and some experiments have shown contrasting results in the affinities of these mutants with H3 peptides. For example, Perlman *et al*. showed that the insertion mutation abolished the interaction between YEATS and H3K27ac, ^12^ whereas Wan *et al*., reported a slightly lower binding affinity of the insertion mutant with H3K27ac than wild type YEATS. ^15^ Moreover, recent findings using isothermal calorimetry (ITC) experiments reported similar binding affinities of wild-type and the deletion mutant for H3K9ac but with different thermodynamics.^16^ Interestingly, structural studies have shown that the overall structure of YEATS remains largely unchanged across mutations, despite the lengthening and shortening of the L8 loop upon insertion or deletion (Figure 1).^16^ Therefore, the mechanisms underlying these disparate binding affinities and how they are affected by mutations remain an open question.

To resolve these differing experimental results and understand the molecular-level effects of mutations on the binding of YEATS to acetylated histones, we performed multiple microsecond-long molecular dynamics (MD) simulations. Here, we have focused on how conformational and thermodynamic differences induced by mutations affect the binding of H3K9ac and H3K27ac peptides. MD simulations revealed that the binding of H3K9ac with

YEATS is enthalpically more favorable than H3K27ac and that residues beyond the acetylation recognition signature (−ARKacS−) in these peptides interacted dynamically with the YEATS domains. Hence, in the full-length nucleosome, only partial exposure of residues is likely required for recognition of acetylation marks on histone tails by YEATS. We also noted that H3K9ac binding highly restricts the conformational space of the deletion mutant in comparison to the wild-type and insertion mutant. Simulations also show favorable binding of H3K9ac peptide to the insertion mutant, in contrast to some previous experimental studies which show no binding, but with altered thermodynamics relative to the wild-type. ^12,16^ In addition, in the context of the nucleosome, the accessibility of histone-acetylation sites (K9, K27) and other positively charged residues is masked by their strong interactions with DNA.^17–19^ Therefore, the specificity of epigenetic modifications is also likely controlled by higher-order interactions of histone tails with nucleosomal DNA, which is not captured by our simulation study and previous experiments.

## Materials and Methods

### Structure Models

The structure of the peptide-free (apo) wild-type YEATS domain was obtained from the Protein Data Bank (PDB) entry 5J9S, ^11^ and the deletion and insertion mutants were from entries 7B10^16^ and 7B0T^16^ (Figure 1). The peptide bound state of the wild-type YEATS domain was from PDB entry 5J9S^11^ and was used for peptide-bound simulations by increasing the peptide length using MODELLER^20^ to reproduce the 11 residue histone H3 tail peptides acetylated at K9 and K27 used for ITC experiments in previous studies^12,21^. Further, structures from 7B10^16^ and 7B0T^11^ were aligned with 5J9S to generate bound states of the YEATS domain insertion and deletion mutants. The peptide sequences used for H3K9ac and H3K27ac were (1-11H3)ARTKQTAR(Kac)ST and (19-29H3)QLATKAAR(Kac)SA, respectively. Both the peptides share a common −ARKacS− motif. We divided the peptide sequences into two regions: a consensus (−ARKacS−) and non-consensus (H3K9ac(ARTKQT) and H3K27ac(QLATKA)) motifs.

### MD simulations

All simulations were conducted using the AMBER software suite with the CUDA-enabled PMEMD engine, version (v18). ^22,23^ Details of all systems are systems are provided in (Table S1). The Amber 14SB force-field parameters along with OPC water models were used, along with previously derived paramaters for acetylated lysines.^24–26^ A 4 fs time-step was used in conjunction with SHAKE and hydrogen mass repartitioning for all simulations.^27,28^ Although in theory mass repartitioning and a longer timestep may alter the observed kinetics and simulation stability, several studies have shown that this repartitioning of mass to hydrogens from heavy atoms, along with the longer 4 fs timestep, has minimal influence on long-timescale dynamics and conformational sampling of diverse all-atom systems ^27,29,30^ All systems were first energy minimized for 5000 steps with a solute harmonic restraint of 10 kcal/mol/Å^2^, followed by 5000 steps without restraints. Equilibration was performed in two steps including 100 ps of constant volume simulations where the temperature was gradually heated from 10 to 310 K. Then, the heavy atom restraints were gradually released over 500 ps of NPT simulations. The pressure was controlled via a Monte Carlo barostat with a target pressure of 1 atm and a relaxation time of 3.0 ps^*−*1^.^31^ Production runs were performed at 310 K using a Langevin thermostat.^32,33^ We performed ten, ^34,35^ 1 µs simulations per system in the NPT ensemble, accumulating 90 µs of sampling across all the three systems. Trajectories were recorded every 10 ps and visualized using VMD^36^ and PyMol. ^37^ Trajectories analyses was performed on the last 750 ns of the simulations, allowing for 250 ns of equilibration using CPPTRAJ^38^ and in-house Python scripts.

### Simulation analyses

Root mean square fluctuation (RMSF) analysis was performed on the apo and peptide bound (H3K9ac and H3K27ac) states of wild-type and mutant YEATS domain. Translations and rotations were removed by least-squares fitting the backbone of the YEATS domains to the initial simulation configurations, and RMSFs were computed on the Cα atoms. Reported RMSFs are the average of all 10 simulations with the standard error of the mean obtained from 10 samples for each system. The Kullback-Leibler divergence of the backbone and side-chain dihedrals was evaluated with the mutinf package to compare conformational ensembles across the systems^39,40^. Contact analysis between YEATS and histone peptides was performed using MDanalysis ^41^ and in-house Python scripts, where contacts were defined as between heavy atoms of histone peptides residues that were within 4.5 Å of YEATS heavy atoms. Interaction energies between the YEATS and acetylated peptides were determined via an MM-GBSA (Molecular Mechanics Generalized Born Surface Area) analysis with igb=8 and a salt concentration of 0.15 M^42,43^ with the MMPBSA.py program. ^42^ Error bars represent the standard error of the mean, with a decorrelation time of 20 ns that is based on a statistical inefficiency test of MM/GBSA values.^44^ Further, to evaluate the conformational entropies of the bound and unbound H3K9ac and H3K7ac peptides, maximum information spanning tree algorithm (MIST) analyses was performed using the PDB2ENTROPY program.^45^ These analyses were performed on the equilibrated portions of trajectories, and forward averaging was performed using a window size of 75 ns to gauge convergence of the conformational entropies.

Principal component analysis (PCA) was used to extract slow and functionally important motions of YEATS domains in apo and peptide bound states from the MD trajectories. First, the covariance matrix of atomic positions of Cα atoms of wild-type and mutant YEATS domains was built, then the eigenvectors of this matrix (also known as principal components) were computed to describe the concerted motion of the system. ^46^ In this study, PCA was performed using GROMACS-2016.3^47^ to investigate the similarities and differences in the conformational ensembles of wild-type and mutant YEATS domain.

A generalized correlation analysis was performed to determine how the motions of individual atoms were correlated with each other. Generalized correlation analysis is a mutual information based method for computing correlations between the Cartesian coordinates of atoms in MD simulations.^32,48^ Unlike a standard mutual information metric, correlations are scaled to an interval of zero to one, in analogy with the more widely used Pearson correlation coefficient. Unlike Pearson correlations, generalized correlation captures linear and non-linear correlations to describe coupled motions. Here, we have used the g correlation program to compute generalized correlation based on the YEATS domain Cα atoms.

## Results

### Mutations and Peptide Binding Alter Loop Dynamics

To assess how mutations and substrate binding affect YEATS domain dynamics, a series of MD simulations were performed. For the wild type, insertion, and deletion mutations, 10 independent 1 µs simulations were run with no substrate bound (Table S1). Additional sets of 10x1 µs simulations were performed for systems with H3K9ac and H3K27ac peptides bound for each protein to determine the disparate effects these two substrates have on system structures, dynamics, and thermodynamics. In each simulation, the YEATS domain maintained a well-folded state with minimal overall structural changes along with a marginally increased mobility of Cα atoms in the regions containing residues 2-30, 45-60, 65-80 in the apo forms of the mutants (Figures 2, S1, S2, and S3). An analysis of protein fluctuations pinpointed two regions of the YEATS domain in which mutations or peptide binding altered RMSF values by more than the standard error: loop 1 and loop 8. Root-mean-square fluctuation (RMSF) analysis of protein Cα atoms showed that the majority of each protein had minimal dynamics, with RMSF values of 1.0-1.5 Å (Figure 2). In loop-8 there were significant differences in RMSF values between the wild-type, deletion, and insertion mutants, regardless of the peptide binding state. In the wild-type, fluctuations were on the order of 2.0 Å (Figure 2a). These fluctuations were reduced in the deletion mutant to ∼1.5 Å, but they were increased to ∼4.0 Å in the insertion mutant (Figures 2b & 2c). This dramatic increase was due to the increased length of this loop in the insertion mutant, whereas in the deletion mutant the loop was shorter and thus restricted its range of motion.

**Figure 2:**
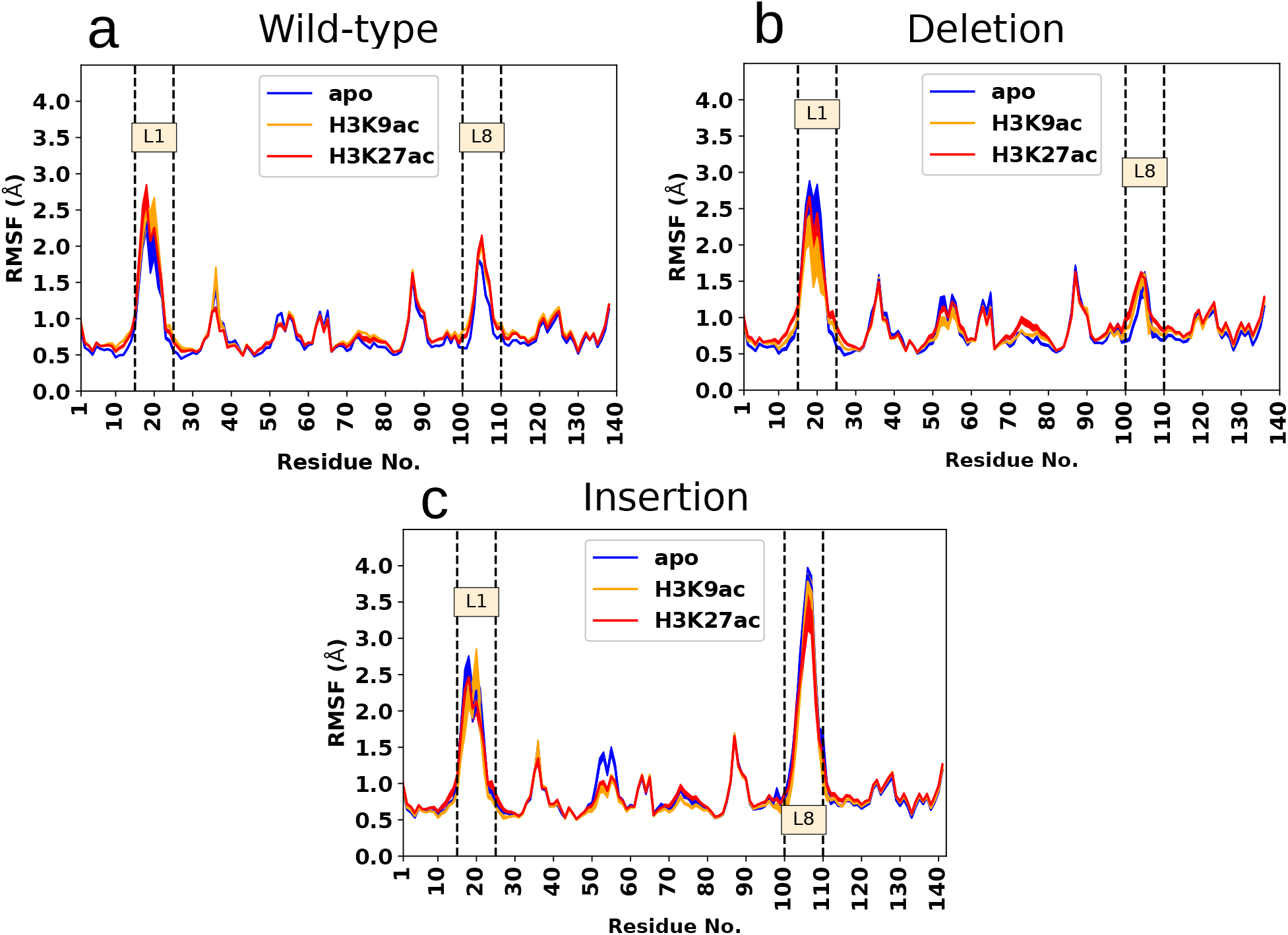
Root mean square fluctuation of Cα atoms in wild-type and mutant YEATS domains with respect to their initial conformations. a) wild-type, b) deletion, and c) insertion. Error bars represent the standard errors over 10 replicate simulations. For a similar plot with standard deviations, see Figure S1.

To probe the effects of mutations and peptide binding on YEATS domain internal dynamics, the distributions of backbone and side-chain dihedral motions were compared between systems with Kullback-Leibler divergence (KLD) calculations. In the apo state, these calculations show that the dynamics are most perturbed by the deletion mutation, which created high KLD values throughout the protein relative to both the wild-type and insertion systems (Figure S4). Meanwhile, the KLD values upon substrate binding were highest upon H3K9ac binding to the wild-type and deletion mutants (Figure S5). To understand which regions of the YEATS domain were most affected by peptide binding, the relative KLD values were mapped onto the protein structure (Figure S6). This showed that binding of H3K9ac introduced long-range allosteric changes in wild-type and deletion mutants, with only minimal effects and an absence of long-range perturbations in the internal dynamics of the insertion mutant. However, for H3K27ac binding the effects on the internal dynamics of YEATS was restricted to the region near the acetylated binding pocket.

Mutational studies have shown that histone peptide recognition is highly dependent on Tyr74, which is located above the acetylated lysine binding pocket.^11^ To monitor the conformation of Tyr74, the distributions of the two sidechain dihedrals, χ1 and χ2, were computed. In the apo wild-type YEATS, Tyr74 sampled a dynamic equilibrium between stacked and unstacked states, with χ1 sampling four states and χ2 independently sampling two, for a total of eight energy wells (Figure 3). Upon peptide binding there was a dramatic shift in these distributions. Binding of the H3K27ac peptide largely restricted χ1 to the well around 100^*°*^, with a similar sampling of χ2. In contrast, the binding of H3K9ac to the wild-type shifted the distribution of χ2 towards positive values, where the Tyr74 hydroxyl group was anti-parallel to the peptide’s acetylated lysine. The Tyr74 conformations were largely unaffected by the insertion mutant, with the apo conformation sampling multiple energy wells which were restricted by peptide binding (Figure 3c). However, the deletion mutant created a dramatic shift in the distributions of the χ1 and χ2 angles. In particular, the χ1 angles were primarily centered around ±180^*°*^ in the apo state, which was maintained in the H3K27ac bound state with similar distributions of both stacked and unstacked Tyr74 conformations. In contrast, upon binding H3K9ac, Tyr74’s χ1 shifts to ∼0 ^*°*^ and χ2 is restricted to ∼-100^*°*^, showing a sampling of multiple stacked conformations with the absence of unstacked conformation

**Figure 3:**
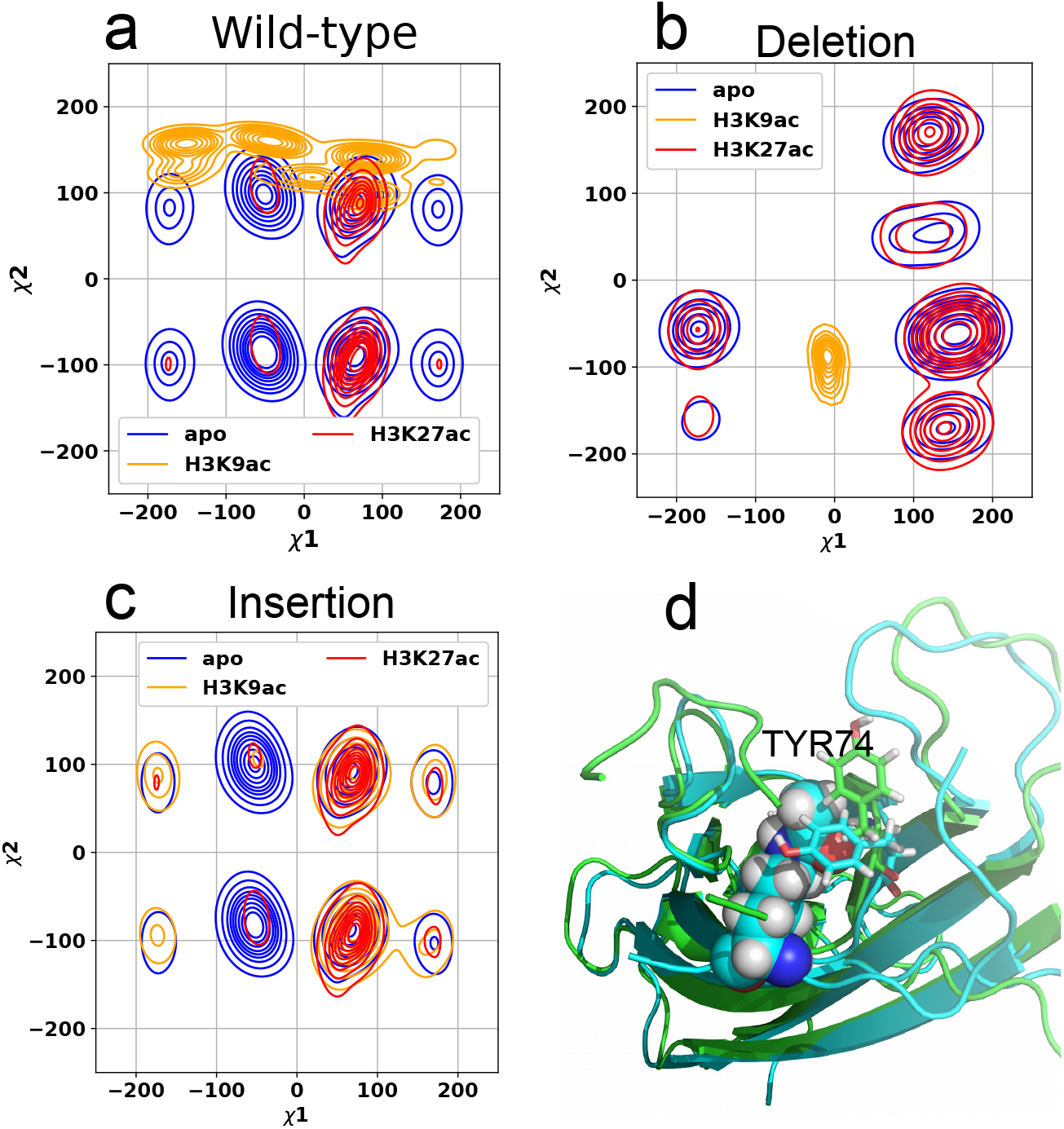
Distribution of side-chain torsional angles (*χ*1, *χ*2) of Tyr74 representing different conformational states of Tyr74 in apo, H3K27ac and H3K9ac bound wild-type, insertion and deletion mutant. a) wild-type YEATS, b)deletion-mutant, and c) insertion-mutant. d) Snapshot representing stacked (cyan) and unstacked (green) conformation of Tyr74 (shown in stick representation).

### Peptide Residues Beyond the Consensus Motif and YEATS Mutations Alter Binding Thermodynamics

To estimate the binding energies of the H3K9ac and H3K27 peptides with wild-type and mutant YEATS domains (ΔE), a series of MM/GBSA analyses were performed (Table 1). We note that MM/GBSA methods involve several approximations, including mean-field solvent approximation and exclusion of configurational entropy, and therefore care must be taken while interpreting the results, which are largely enthalpic in nature. To partially account for these shortcomings, we have also estimated the change in conformational entropy of each peptide upon binding to each YEATS domain, along with the associated change to the free energy -TΔS, using MIST analyses (Table 1 and Figure S8). When added to MM/GBSA energies, these values provide a more complete, although still qualitative, binding free energy estimate (ΔG).

**Table 1:** MM/GBSA derived binding energies, peptide conformational entropies, and total binding energies of H3K9ac and H3K27ac with wild-type and mutant YEATS domains. All values are given in kcal/mol with standard error of the mean calculated across 10 simulations. Binding energies presented here should be interpreted comparatively, rather than absolute quantities.

| System | Peptide | MM/GBSA Energy ($\Delta E$) | Peptide Conformational Energy ($-T\Delta S$) | Total Binding Energy ($\Delta G$) |
| --- | --- | --- | --- | --- |
| Wild-type | H3K9ac | $-43.94 \pm 1.05$ | $8.22 \pm 0.74$ | $-35.72 \pm 1.79$ |
| | H3K27ac | $-40.27 \pm 0.28$ | $5.60 \pm 0.30$ | $-34.67 \pm 0.58$ |
| Deletion Mutant | H3K9ac | $-45.58 \pm 0.72$ | $7.98 \pm 0.63$ | $-37.55 \pm 1.35$ |
| | H3K27ac | $-40.26 \pm 0.41$ | $6.49 \pm 0.30$ | $-33.51 \pm 0.71$ |
| Insertion Mutant | H3K9ac | $-49.38 \pm 1.01$ | $11.00 \pm 0.84$ | $-38.38 \pm 1.85$ |
| | H3K27ac | $-41.94 \pm 0.45$ | $7.07 \pm 0.55$ | $-34.87 \pm 1.00$ |

The results of these thermodynamic analyses showed peptide identity and YEATS mutations affected the interplay between the binding energies and entropies of peptides to YEATS domains. For example, in the wildtype domains, H3K9ac was slightly more energetically favored to bind than H3K27ac, with a difference in binding energies (which we denote as ΔΔE) of -3.67±1.09 kcal/mol. This energy difference was offset by an increased entropic penalty of H3K9ac binding (-TΔΔS=2.62±0.84 kcal/mol) which resulted in a binding free energy difference between the peptides which was smaller than the error bars (ΔΔG=-1.05±1.89 kcal/mol). This trend of more favorable ΔE values for H3K9ac over H3K27ac, along with higher entropic penalties, was present in both the deletion and insertion mutants. However, the degree of these differences between peptides was different in both mutants. In the deletion mutant, the ΔΔE between peptides expanded to -5.32±0.83 kcal/mol, whereas the -TΔΔS term was reduced to 1.49±0.70 kcal/mol, for a -ΔΔG of 4.04±1.53 kcal/mol, thus favoring H3K9ac binding over H3K27ac. In the insertion mutant, both the entropic and enthalpic differences were magnified between peptides (ΔΔE=7.44±1.10 kcal/mol and -TΔΔS of 3.93±1.00 kcal/mol), with a net result of a similar ΔΔG of -3.51±2.1 kcal/mol. H3K9ac and H3K27ac have similar residues in the consensus motif (−ARKacS−) but differ mainly in the N-terminal region with H3K9ac being more positively charged due to the presence of an additional arginine in the N-terminus. To understand the role of the peptide’s primary sequence on binding thermodynamics, we analyzed the per-residue contributions in the binding energies, which revealed that the presence of Arg2 can provide modest energetic stabilization of ∼1.2 kcal/mol in H3K9ac as opposed to the negligible contribution of Leu2 in H3K27ac at the same position (Figures 4a, and 4b). Further, additional residues in H3K27ac (Ala6, Lys5) and H3K9ac (Thr6, Gln5), which are adjacent to the consensus motif, provided higher energetic stabilization in the insertion mutant than the wild-type and deletion. These difference resulted in a higher number of contacts and more favorable binding energies of both peptides to the insertion mutant (Figure 4c).

**Figure 4:**
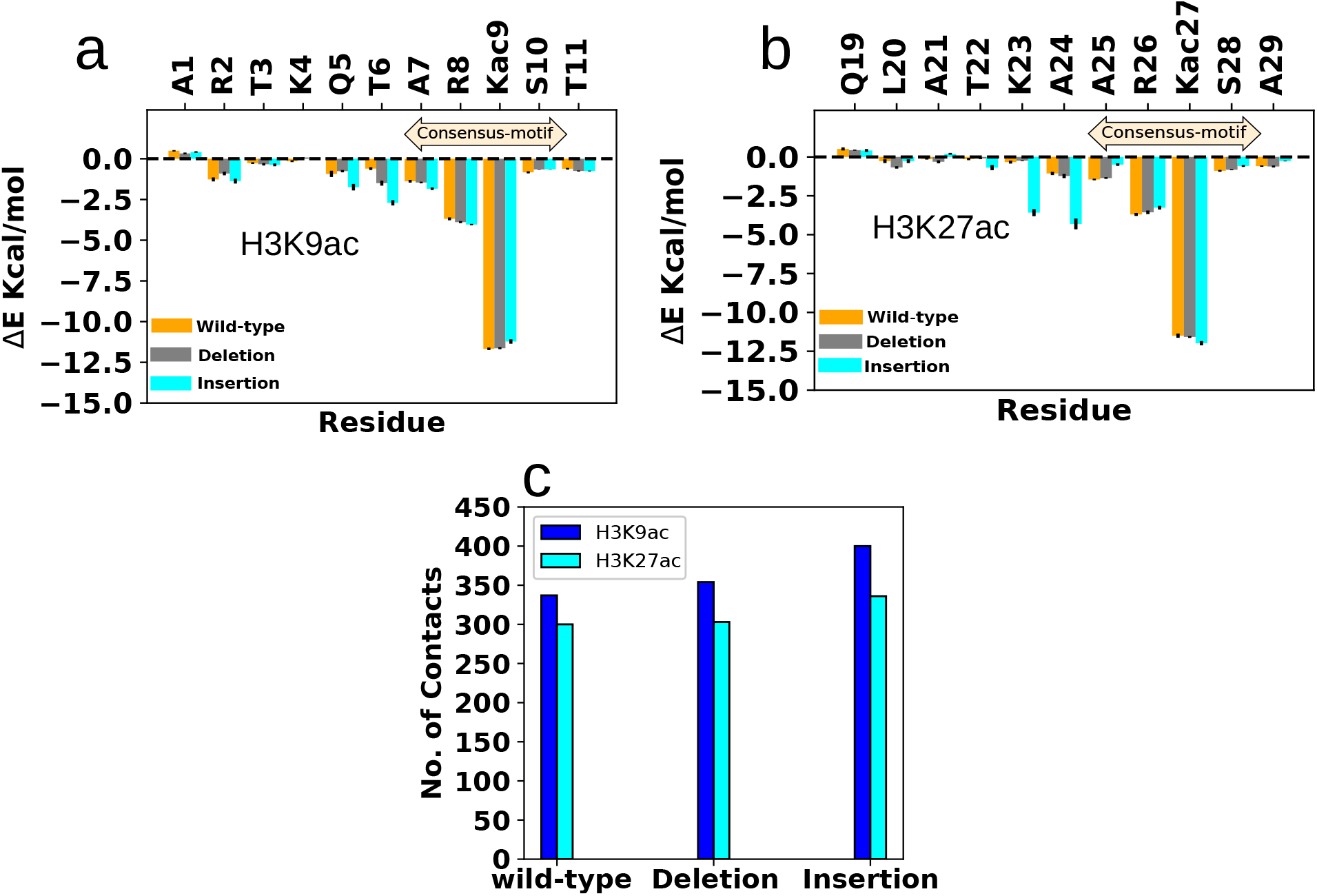
MM/GBSA derived per-residue binding energies of peptides in complex with wild-type and mutant YEATS domains a) H3K9ac, b) H3K27ac, and c) Contacts formed between histone peptides (H3K9ac and H3K27) heavy atoms and wild-type and mutant YEATS domain within a distance cut-off of 4.5 Å.

In addition to the previously noted differences in YEATS domain flexibilities, there were significant differences in the flexibility of the H3K9ac and H3K27ac peptides in the bound state. To quantify this, we computed RMSFs of the peptides relative to the YEATS domain in all systems, which is an indirect measurement to assess the conformational freedom of histone peptides upon binding (Figure S7). The N-terminal region of the H3K9ac and H3K27ac in the YEATS domain had large fluctuations, with RMSF values in the range of 410 Å and 5-12 Å, respectively, with consistently higher RMSFs in H3K27ac. The C-terminal region of the H3K27ac fluctuated in the range of 3-5 Å, whereas H3K9ac fluctuated in the range of 2-3 Å. We also observed a marginal increase in the flexibility of the C-terminal region of H3K27ac when bound to the deletion mutant. These RMSF differences highlight the role of peptide chemistry and mutations in modulating the peptide dynamics in the YEATS complex.

### Peptide binding alters large-scale YEATS domain motions

To dissect the large-scale motions and characterize the essential dynamics of YEATS domains, principal component analysis (PCA) was performed on the conformational region conserved across the wild-type and mutant apo and holo simulations (Figure 5). Conformational ensembles from all simulations were combined in the PCA to ensures that an identical eigenbases were used in comparisons across systems. The first two projections, PC1 and PC2, represent 37% and 26% of the total sampling across all simulations and are the only two projections with more than 10% of the total sampling variance (Figure S9). PC1 and PC2 represent a rocking and twisting motion of the YEATS domain, respectively (Figure 5d, 5e). Projections of the unbound wild-type simulations on PC1 ranged between -1.0 nm and 1.0 nm, whereas PC1 projections from both mutants showed a marginal increase in a positive direction (PC1 -1.0 nm to 1.5 nm), indicating the absence of significant conformational changes upon mutations. Peptide binding only slightly restricted the motion of the YEATS domain in the PC1 direction, confirming the similar amount of conformational dynamics relative to the apo-states, which is in agreement with the RMSF analysis. In the PC2 space we noted a remarkable shift in the H3K9ac bound deletion mutant where the majority of the conformations were in the positive direction as compared to the other system. To determine the degree of conformational subspace overlap between PC1 and PC2 of the H3K9ac bound deletion mutant with all-other systems, the inner-products of the PC1 and PC2 eigenvectors from all systems with the PC1 and PC2 eigenvectors from just the H3K9ac deletion mutant were calculated. Comparison of PC1s gave an inner-product of 0.92, indicating nearly identical motions. In contrast, the inner-product of the PC2s was only 0.031. Inspection of the H3K9ac deletion PC2 showed a motion that was nearly orthogonal to the outward twisting motion present in all other systems with an inward twisting motion.

**Figure 5:**
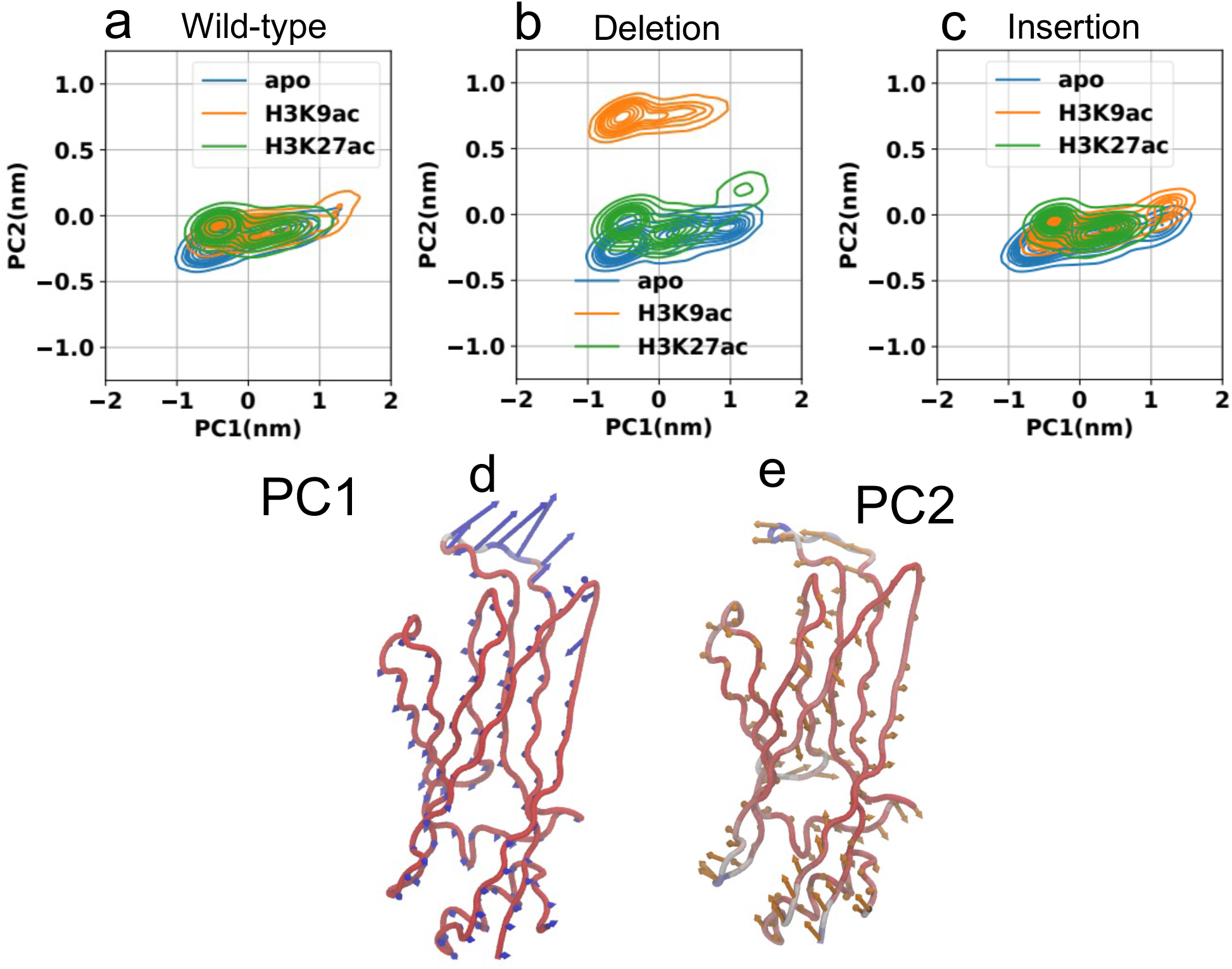
Conformational restrictions in peptide bound states of YEATS domain displayed by the motion along PC1 and PC2. PC2 suggested a inward twisting motion in deletion mutant (b) upon the binding of H3K9ac. PC1 suggested only minor variation in the conformational dynamics of wild-type (a) and insertion mutant(c). Procupine plots showing the direction of the motion of Cα atoms in YEATS domain along PC1 and PC2 representing the rocking and twisting motions.

### Mutations weaken inter-residue correlations

To understand how mutations modulate inter-residue communication, pairwise generalized correlation coefficients of Cα atoms were computed in the peptide bound and unbound states for each system (Figure 6). The thick lines colored in red perpendicular to the diagonal in the plots represent the short-range correlation between anti-parallel β-strands, while dense irregular blocks show correlations between β-sheets (Figure 6a-c). β-strands and inter-β-sheet regions in wild-type YEATS exhibit a strong correlation, which was reduced upon insertion and deletion of residues in loop-8 (Figure 6a-c). This reduced correlation is likely due to the increased flexibility of mutants relative to the wild-type (Figures S1 and S2), which were highest for the insertion mutant and resulted in decoupling of short-range correlations and alterations of long-range correlations.

**Figure 6:**
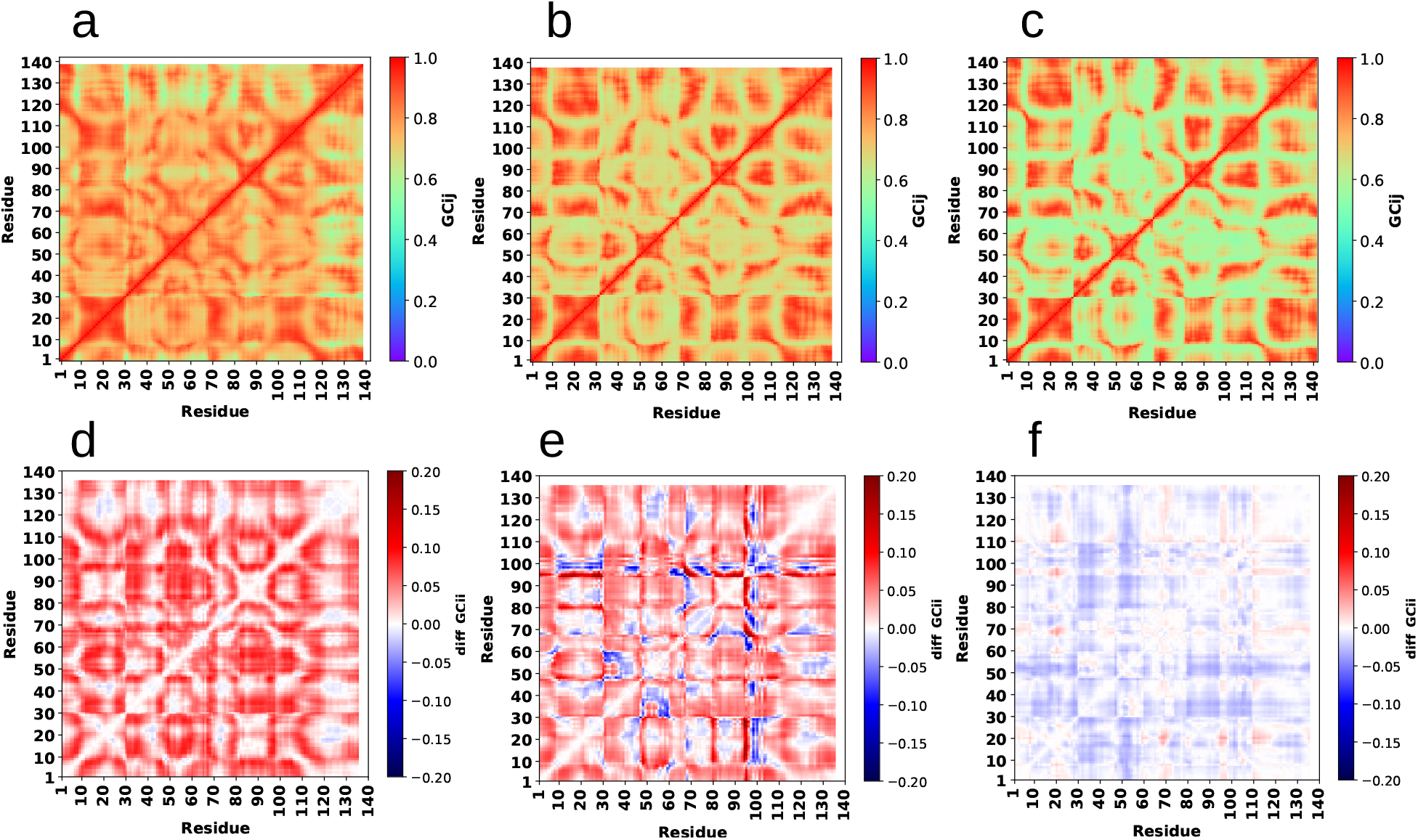
Decreased correlation in apo states of deletion (b) and insertion mutant (c) relative to the wild-type (a). Difference of generalized correlations of H3K9ac bound states to the apo-states of YEATS (d) wild-type (e) deletion mutant and (f) insertion mutant.

To understand how peptide-binding modulates short and long-range correlations, the difference between apo and peptide (H3K9ac and H3K27ac)-bound correlations was calculated (Figure 6d-f & S10). The wild-type YEATS-bound H3K9ac system showed a reduced coupling in both local and distant regions, whereas the dynamics became more correlated between distant regions in the deletion mutant, although there was a net reduction in shortrange correlations. In contrast, the dynamic coupling was largely unaffected by the binding of H3K9ac to the insertion mutant.

H3K27ac binding had a less drastic effect on the inter-residue dynamic correlation in the wild-type and deletion mutant relative to their apo-states. In the deletion mutant, there were no regions in which there was an increase in correlations upon H3K27ac binding and there were several regions in which there was a decrease in correlations (see Figure S10). Moreover, the coupling between structural regions in the insertion mutant remained unaffected upon H3K27ac binding (Figure S10). The changes observed in the correlation pattern across the apo and peptide-bound states of YEATS were highly dependent on the peptide sequence and the type of mutation present in the YEATS.

## Discussion

Mutations in ENL YEATS are linked to altered histone acetylation mark binding abilities.^12,16^ To date, the structural bases of these altered binding mechanisms, and how they influence the binding thermodynamics and specificities of these systems, remains poorly understood, as structural experiments have yet to reveal the binding mechanism of the fulllength peptides used in ITC experiments. This has significantly affected our understanding of how the shift in histone tail acetylation binding preferences in mutant YEATS that are observed in binding assays translates to its altered chromatin occupancy.

Here, we have used extensive MD simulations to discern the differences in the binding dynamics of H3K9ac and H3K27ac with wild-type and mutant ENL YEATS domains. RMSF, KLD, and PCA analyses have revealed differences in wild-type and mutant YEATS domain dynamics in their apo and peptide-bound states. RMSF showed that all apo and bound states have only minor variations in the β-sheet core and that L8 has the largest length and dynamics in the insertion mutant, followed by the wild-type, and then the deletion mutant. Interestingly, in each YEATS domain, peptide binding did not affect the dynamics of L8, which showed a similar level of dynamics relative to their apo forms (Figure 2). However, we observed system-specific changes in other YEATS structural elements. For example, L1 dynamics were marginally reduced in the deletion mutant upon H3K9ac binding, as highlighted by a restriction in the motion along PC1, and an inward twisting motion of the domain was present, as shown in the drastic shift in PC2 (Figure 5). KLD analysis, which measures the difference between internal residue motions, highlighted significant changes in the wild-type YEATS and deletion mutant’s internal dynamics upon H3K9ac binding relative to its apo and H3K27ac bound states, suggesting that H3K9ac induces unique conformational arrangements in the wild-type and deletion mutant (Figure 3). In contrast, the insertion mutant internal dynamics remained largely unaffected by peptide binding. These long-range perturbations in the wild-type and the deletion mutant suggest allosteric changes upon substrate binding, although there have yet to be experimental reports of allosteric communication in YEATS domains. Results of our correlation analyses show that the internal YEATS domains dynamics are highly correlated, making YEATS domains potential candidates for allosteric modulation by small molecules.

Our per-residue energetic calculations showed that binding to YEATS domains is energetically dominated by the −ARKacS− consensus motif present in both peptides. In particular, the arginine in the consensus motif tethers the peptide along the β-sheet by forming a salt-bridge with YEATS’ Asp99 residue (Figure S12). In addition, this analysis also showed that residues beyond −ARKacS− tune the overall affinity of peptides with YEATS domains despite their overall increased dynamic binding (Figure S7). The deletion and insertion mutants have unique interactions with these these residues, which alters the binding thermodynamic and specificity for H3K9ac and H3K27ac. For example, static modelling and X-ray derived structures have led to the speculation that alteration of L8 in the insertion creates increased steric hindrance with the N-terminals of H3K9ac and H3K27ac peptides, and that these differences result in increased specificity for H3K27ac over H3K9ac.^16^ Our per-residue energetic analysis showed that the insertion mutant can interact more favorably with the N-terminal residues K23 and A24 of H3K27ac compared to the deletion and wildtype due to the longer L8 loop, an observation which is consistent with a crystal structure of the H3K27ac bound insertion mutant.^49^

For each YEATS domain and peptide, simulations showed that binding was enthalpically favored and entropically disfavored. In all cases H3K9ac had a stronger binding enthalpy than H3K27ac, which is partially due to the presence of an arginine in the non-consensus residues of H3K9ac that reliably created an additional ∼1 kcal/mol of binding enthalpy and enabled H3K9ac to interact with the L1 loop (Figures 4, 7, and S11). However, YEATS mutations affected the balance of these two terms such that the binding free energy preference for H3K9ac was altered relative to H3K27ac. In the deletion mutant, we noted an increased enthalpic preference, along with a decreased entropic difference in binding of the two peptides relative to the wild-type, which created a net favorable binding of H3K9ac over H3K27ac. This is in line with experiments that have shown the deletion mutant having different thermodynamic signatures for each peptide’s binding relative to the wild-type along with a five-times greater preference for H3K9ac over H27Kac. ^12,15,16^ This is also agreement with work by Ni *et al*., who reported a favorable enthalpic contribution in H3K9ac binding with the deletion mutant.^16^

**Figure 7:**
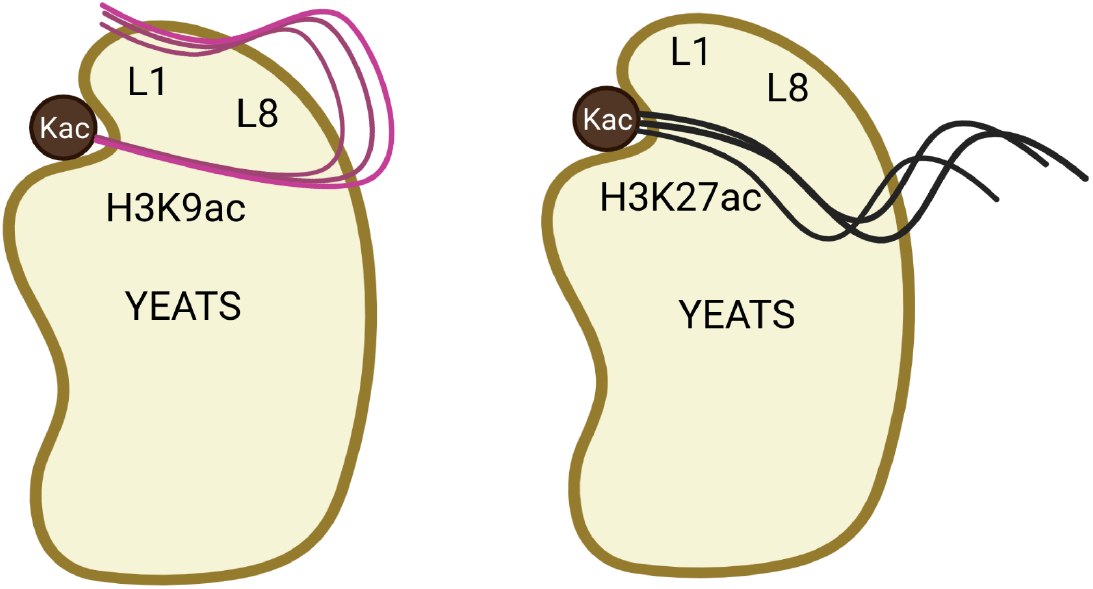
Representation showing the overall differences in the binding of H3K9ac and H3K27ac to YEATS domain. Non-consensus motif of H3K9ac binds with L1 loop, whereas H3K27ac binds more dynamically to YEATS domain.

The energetic analysis also showed that in the insertion mutant there was a larger enthalpic preference and a significantly greater entropic penalty for H3K9ac binding over H3K27ac. The ability of the insertion mutant to bind to H3K27ac is in accord with the reported binding of the insertion YEATS to a H3K27ac peptide. ^15^ However, there is a discrepancy in the favorable binding free energy to H3K9ac, as experiments have suggested that insertion mutants do not strongly bind this substrate. We reiterate that the thermodynamic analyses performed here are qualitative and include several approximations. Although we have attempted to overcome some of them with entropic calculations of the peptides, the YEATS domain itself is too large for similar calculations. However, we note that in the insertion mutant L8 is highly mobile (Figure 2), which likely results in a much more significant entropic penalty to the H3K9ac/insertion YEATS domain binding than is captured by our calculations. However, taken with biochemical experiments, our calculations highlight that it is the entropic term, and not the enthalpic term, that appears to impair the binding of peptides, especially H39ac, to the insertion mutant.

It has previously been shown that in the context of full nucleosomes, exposure of histone tails are limited by histone tail/DNA interactions.^17,50,51^ This observation has direct implications for the mechanisms by which chromatin readers such as YEATS recognize PTMs. For example, in the YEATS insertion mutant, it was recently shown that despite the weaker binding affinity to H3K27ac, it can robustly bind to acetylated chromatin. ^15^ In addition, studies on the ENL analog AF9 YEATS domain showed an additive role of acetylation and the engagement of two YEATS domains on the same histone tail in the context of a full nucleosome. ^52^ This study also highlighted that YEATS domains engage primarily with histone tail consensus motifs when they interact with an acetylated nucleosome core particle.^52–54^ Our MD studies also support that only partial exposure of the histone tails may be required for binding to full nucleosomes, as the consensus motif and the residues directly adjacent to it have the largest influence on binding thermodynamics. *In vitro*, engagement of residues non-adjacent to the consensus motif, along with increased L8 length dynamics, may have a reduced effect on peptide selectivity than observed here or in experiments with histone tail mimics, as those residues may be interacting with nucleosomal DNA. This may create effects which are distinct from those studied with histone tail mimics. For example, the insertion and deletion mutant have unique peptide binding properties, but in the full nucleosome their differences may be reduced, which would result in distinct tumorigenesis phenotypes then is expected from experiments with histone tail mimics. Further computational and experimental studies are needed in the context of full-length nucleosomes to understand how mutations linked with diseases modulate the preference of YEATS domains for lysine acetylation marks on histone tails.

## Data and Software availability

Details for all the data generated in this molecular dynamics study are available in the Methods section including PDB codes of input files^11,16^ and simulation protocol. The simulation^22,23^ and visualization software^36,37^ and the force field^24^ used to perform this study are openly available. MD trajectories generated in this study are available on Zenodo at: https://doi.org/10.5281/zenodo.6870475.

## Acknowledgement

The authors thank members of the Wereszczynski Group for valuable discussions concerning this work. This research was supported by the National Institute of General Medical Sciences of the National Institutes of Health Grant R35GM119647 and the National Science Foundation grant CAREER-1552743. The content is solely the responsibility of the authors and does not necessarily represent the official views of the National Institutes of Health.

