## Supporting Information for "Conformational and Thermodynamic Differences Underlying Wild-type and Mutant ENL YEATS Domain Specificity for Epigenetic Marks"

Table S1: Details of all the simulations carried out in this study

| Protein | Peptide | Number of Simulations | Simulation Length | Total Sampling |
|---|---|---|---|---|
| None | H3K9ac | 10 | 1 µs | 10 µs |
| | H3K27ac | 10 | 1 µs | 10 µs |
| Wild-type | None (apo) | 10 | 1 µs | 10 µs |
| | H3K9ac | 10 | 1 µs | 10 µs |
| | H3K27ac | 10 | 1 µs | 10 µs |
| Insertion Mutant | None (apo) | 10 | 1 µs | 10 µs |
| | H3K9ac | 10 | 1 µs | 10 µs |
| | H3K27ac | 10 | 1 µs | 10 µs |
| Deletion Mutant | None (apo) | 10 | 1 µs | 10 µs |
| | H3K9ac | 10 | 1 µs | 10 µs |
| | H3K27ac | 10 | 1 µs | 10 µs |
| | | Total Sampling Time: | | 110 µs |

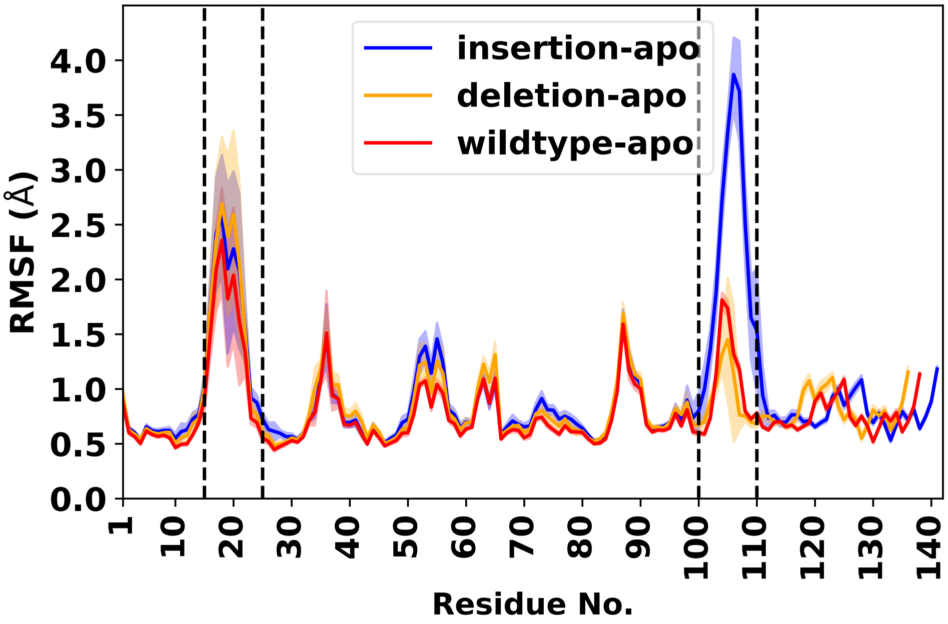


(a)

Figure S1: Root mean square fluctuation of Cα atoms present in apo states of wild-type and mutant yeats domain with respect to their initial conformation. Error bars represent the standard deviations over 10 replicate simulations.

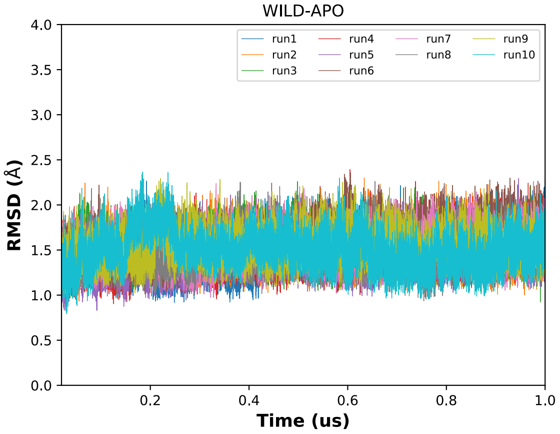


(a)

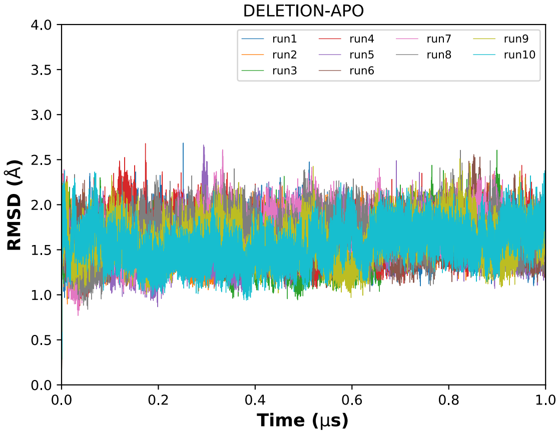


(b)

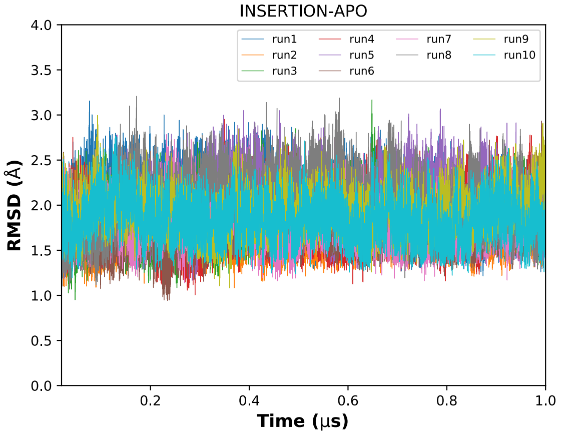


(c)

Figure S2: Root mean square deviation (RMSD) of C$\alpha$ atoms present in YEATS domains. a) wild-type, b) deletion, and c) insertion, respectively

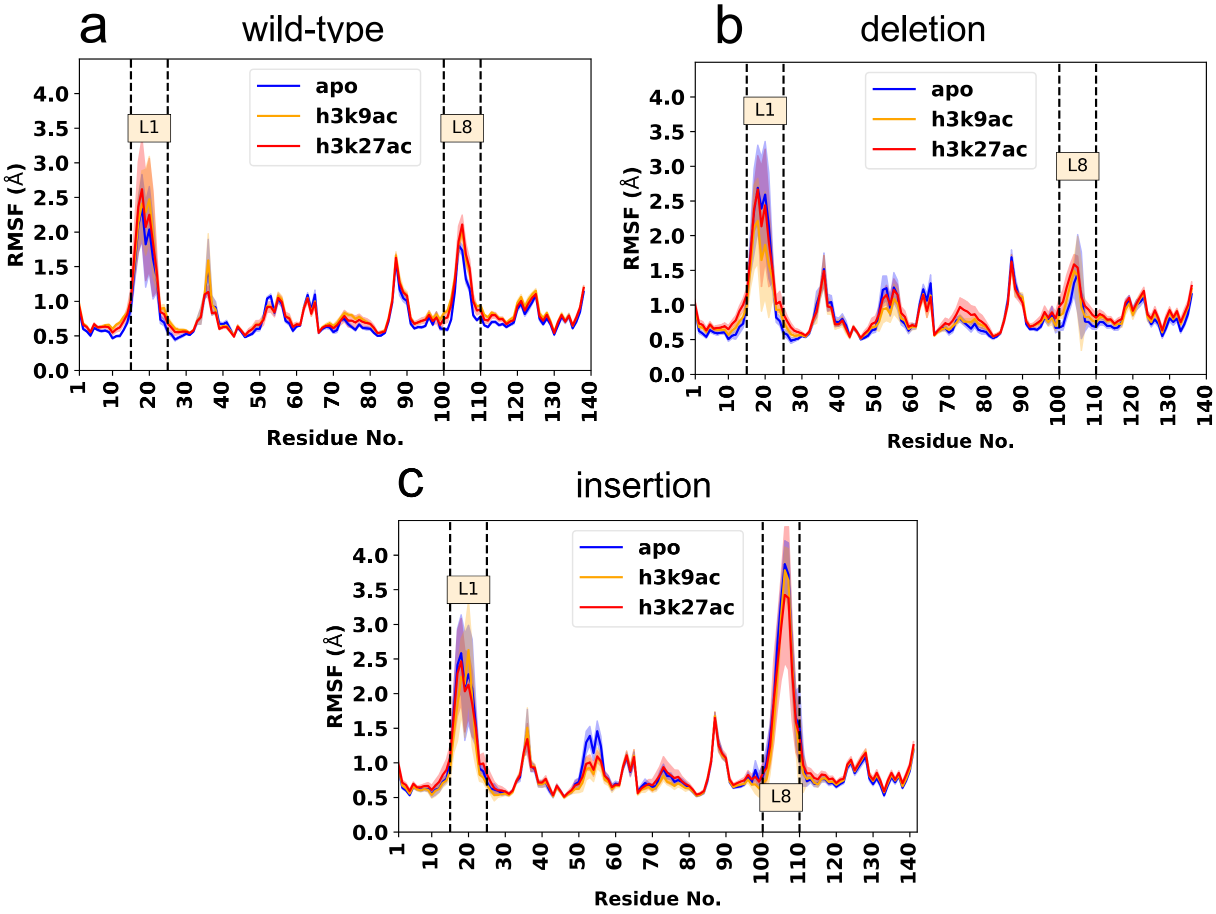


Figure S3: Root mean square fluctuation of Cα atoms present in wild-type and mutant yeats domain with respect to their initial conformation analyzed over 10 replicates. a) wild-type, b) deletion, and c) insertion. Error bars represent the standard deviation over 10 replicate simulations.

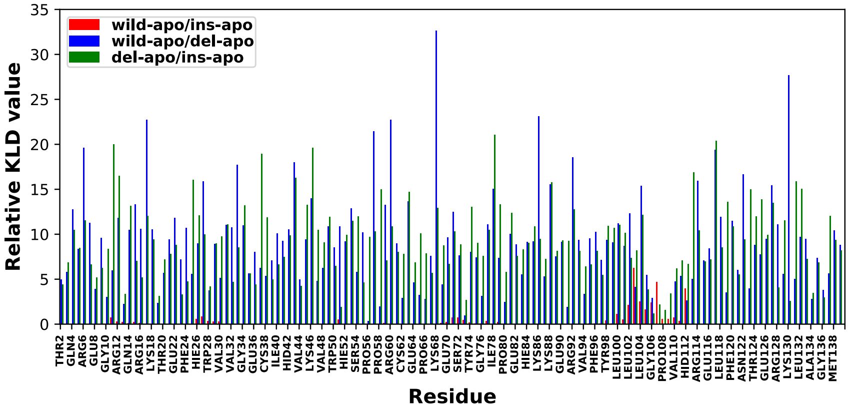


Figure S4: Relative Kullback-Leibler Divergence (KLD) value obtained after comparing the backbone and side-chain torisonal space of wild-type and mutant YEATS.

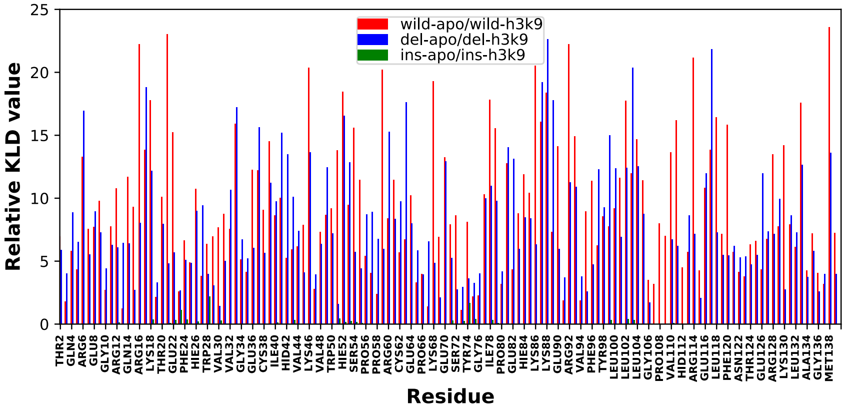


(a)

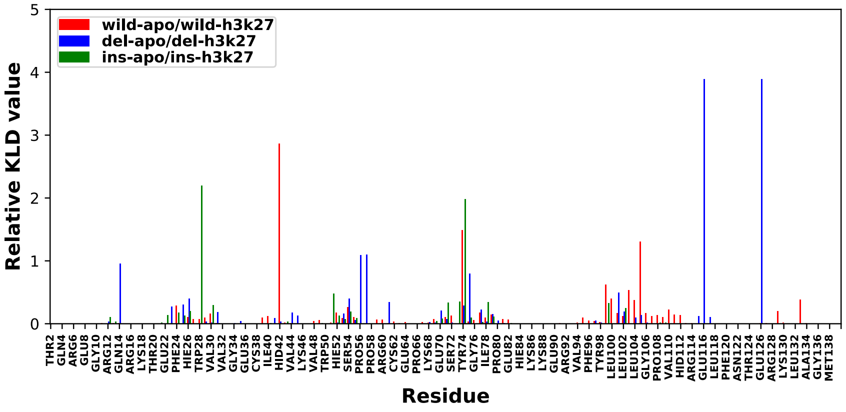


(b)

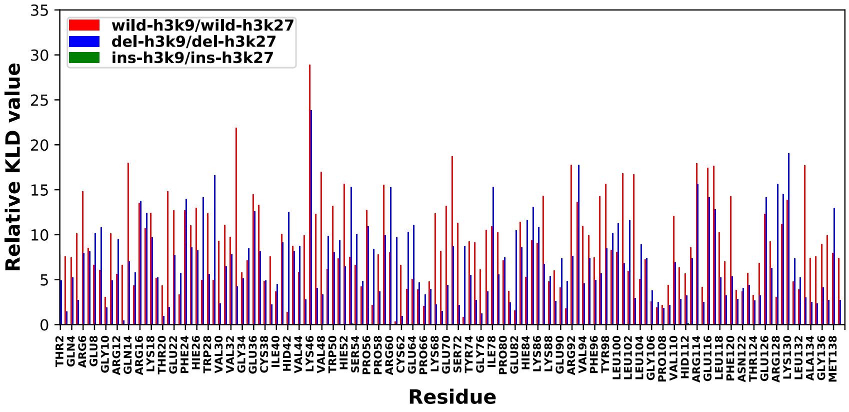


(c)

Figure S5: Relative Kullback-Leibler Divergence (KLD) value obtained from wild-type and mutant YEATS system a) Per-residue KLD of YEATS with apo states as reference ensemble and H3K9ac as target, b)Per-residue KLD values of H3K27ac bound states, c) KLD of H3K27ac relative to H3K9ac

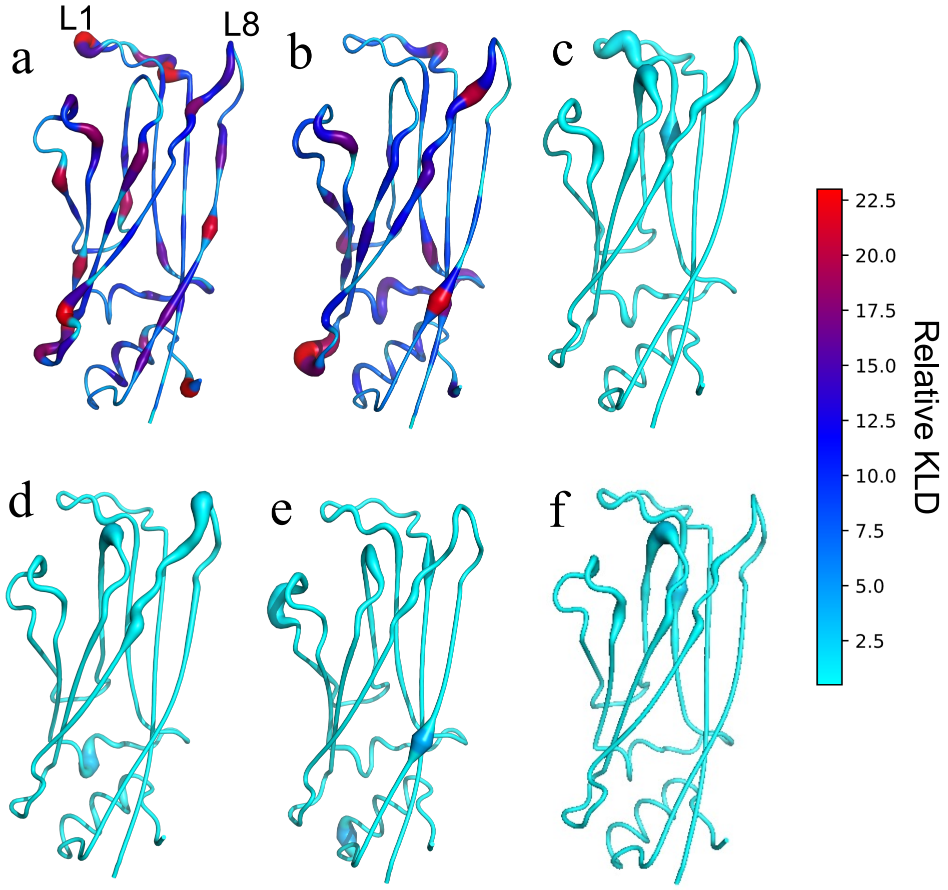


(a)

Figure S6: Overlay of relative KLD values on the YEATS domain compared to their apo states a) wild-type-h3k9ac, b)deletion-h3k9ac, c) insertion-h3k9ac, d) wild-type-h3k27ac, e) deletion-h3k27ac, f) insertion-h3k27ac)

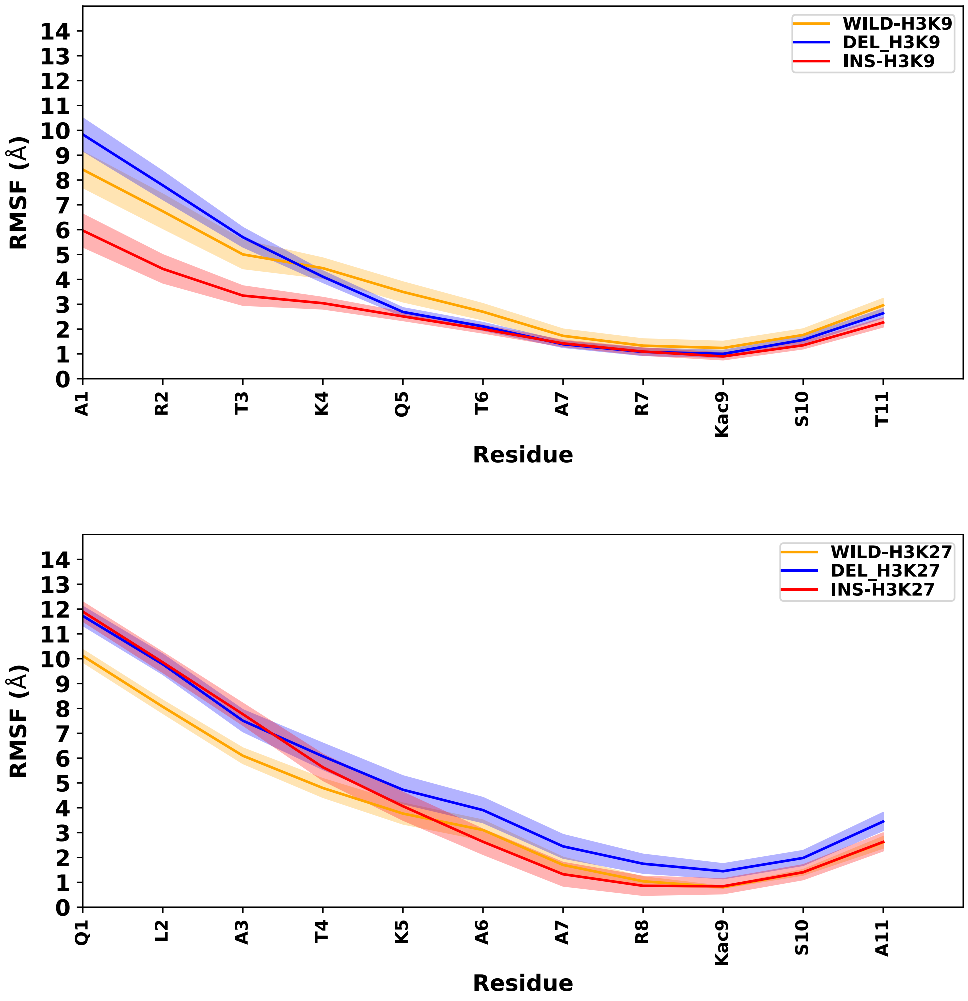


Figure S7: Root-mean square fluctuation of C$\alpha$ atoms of H3K9ac and H3K27ac.

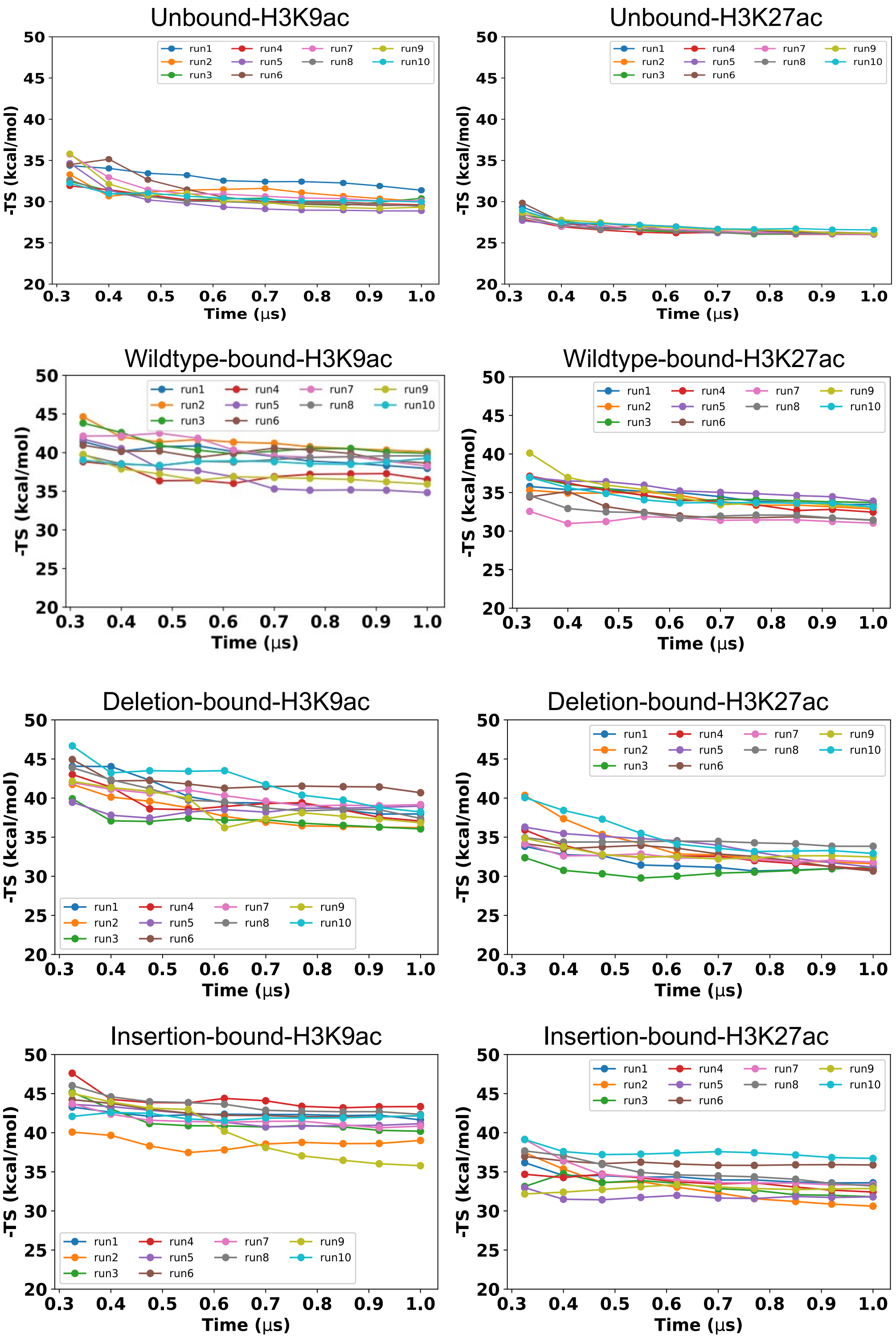


Figure S8: Conformational entropies estimates for unbound and bound H3K9ac and H3K27ac peptides

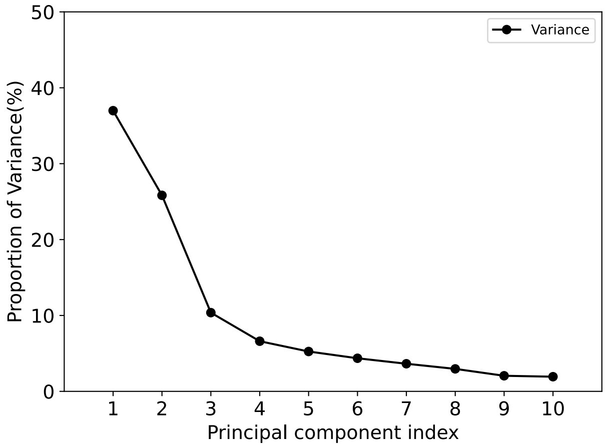


(a)

Figure S9: Proportion of Variance for the first 10 eigenvectors in describing the overall motion of the YEATS domain

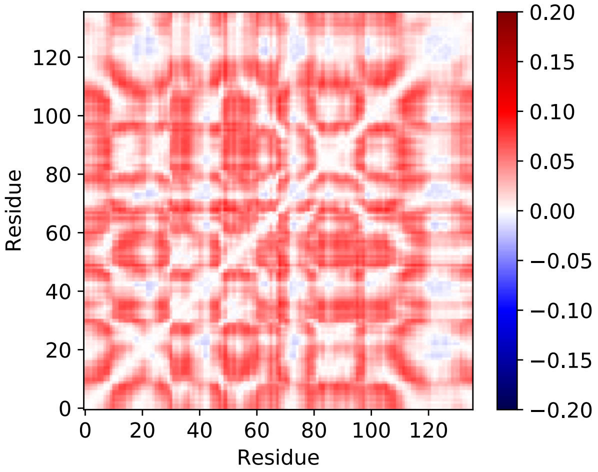


(a)

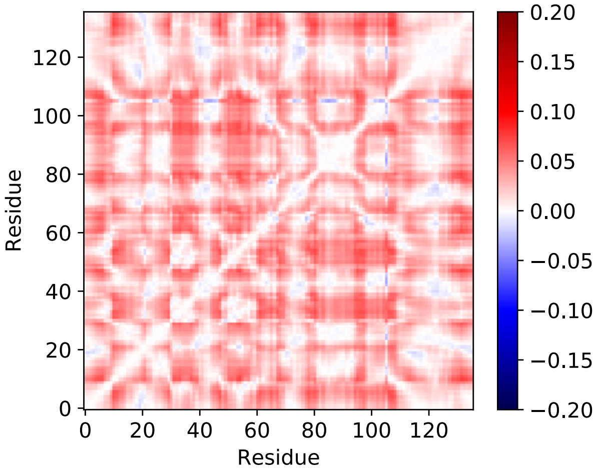


(b)

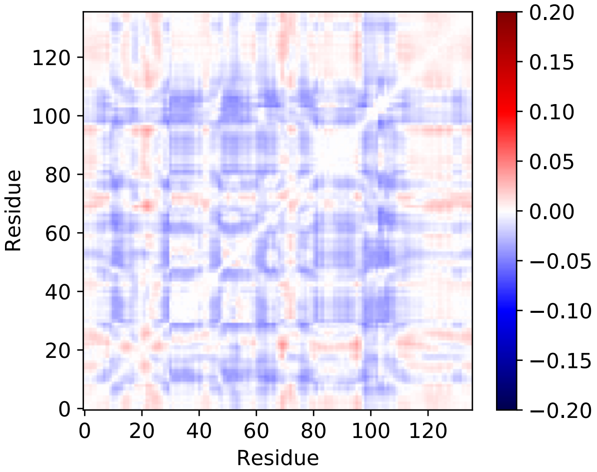


(c)

Figure S10: Difference of generalized correlations between apo and H3K27ac peptide bound state of YEATS domains a) wild-type, b) deletion, and c) insertion, respectively

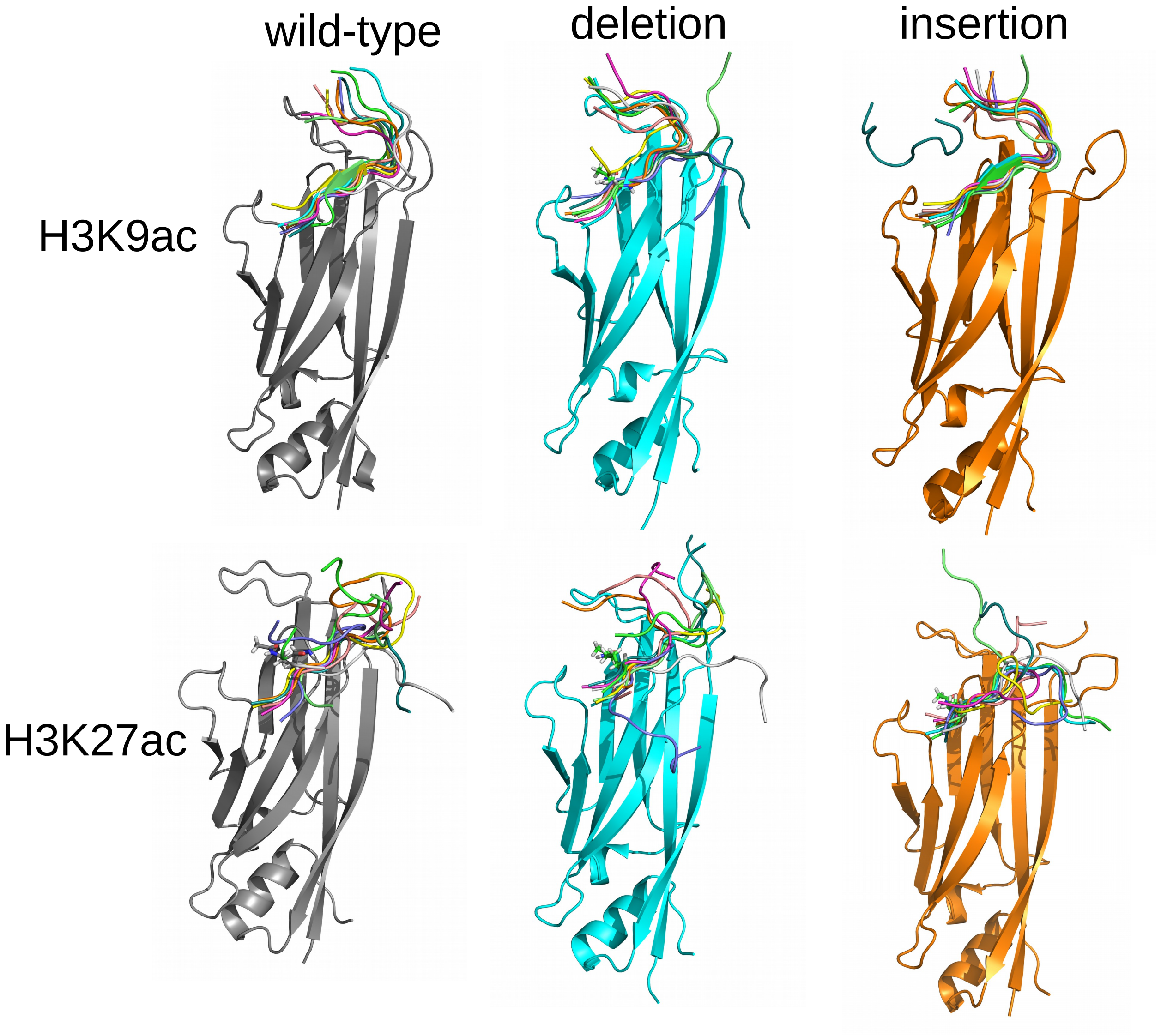


Figure S11: Final conformations of the H3K9ac and H3K27ac peptides bound to wild-type and mutant YEATS domain.

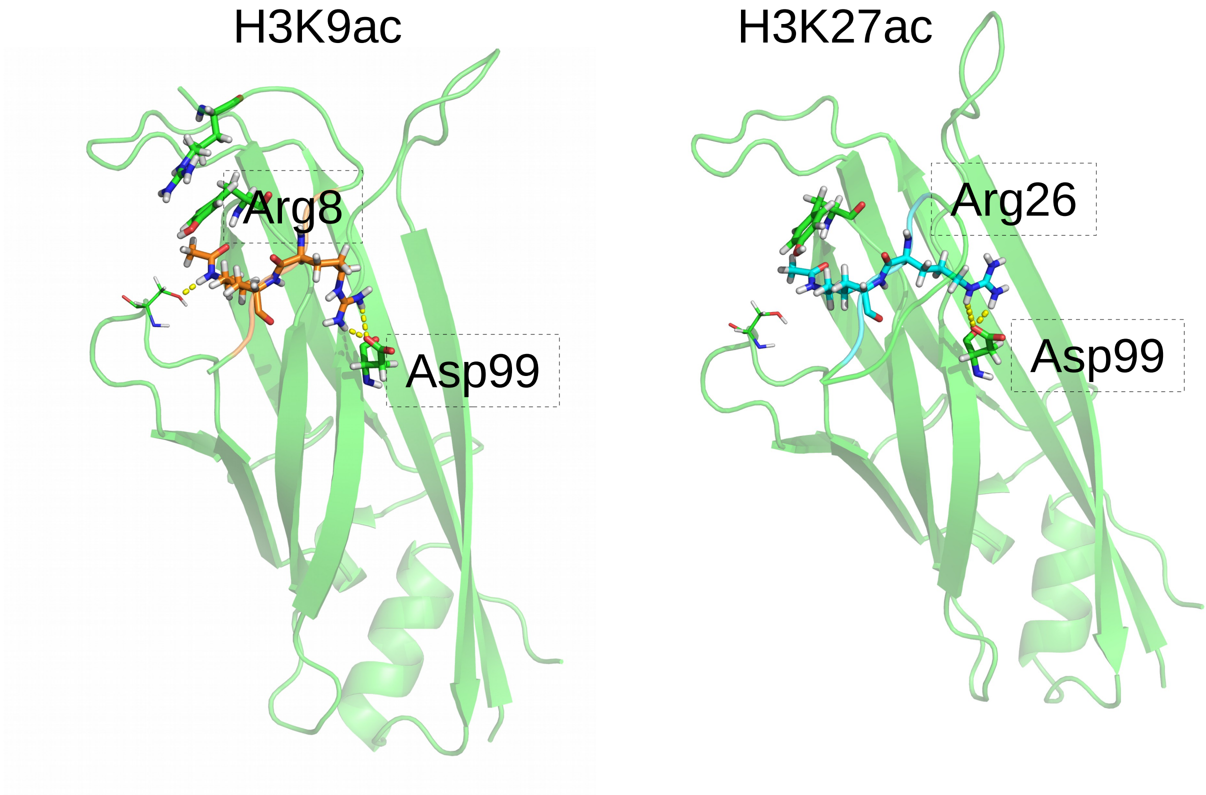


Figure S12: Interaction of arginine of peptide's consensus motif with aspartate of YEATS domain in H3K9ac and H3K27ac system
